# Complex naturalistic stimuli are prioritized by attention when maintained in visual working memory

**DOI:** 10.1101/2020.03.23.004192

**Authors:** Natalia Rutkowska, Łucja Doradzińska, Michał Bola

**Author notes:** **Corresponding author:** Michał Bola, PhD Laboratory of Brain Imaging Nencki Institute of Experimental Biology 3 Pasteur Street 02-093 Warsaw, Poland. **Authors contributions:** conceived study: MB; designed study: NR, ŁD, MB; collected data: NR; analysed data: NR, ŁD, MB; drafted manuscript: NR; revised manuscript: ŁD, MB.

## Abstract

Recent studies suggest that a stimulus actively maintained in working memory (WM) automatically captures visual attention when subsequently perceived. Such a WM-guidance effect has been consistently observed for stimuli defined by simple features, such as colour or orientation, but studies using more complex stimuli provide inconclusive results. Therefore, we investigated whether the WM-guidance effect occurs also for naturalistic stimuli, whose identity is defined by multiple features and relations among them, specifically for faces and houses. The experiment consisted of multiple blocks in which participants (N = 28) either memorized or merely saw (WM or exposure condition, respectively) a template stimulus and then performed several dot-probe trials, with pairs of stimuli (template and control) presented laterally as distractors and followed by a target-asterisk. Evidence for attentional prioritization of the memorized stimuli was found in the reaction-times (RT) analysis, but not in the analysis of the N2pc ERP component, which raises questions concerning the attentional mechanism involved. Further, in an exploratory ERP analysis we found evidence for a very early (100-200 ms post-stimulus) prioritization specific to the memorized faces, which is in line with the sensory recruitment theory of WM.

## Introduction

Contemporary theories of memory emphasize its role in the prospective guidance of perception and action (Nobre and Stokes, 2019). Particularly, the working memory (WM) system is currently recognized as the key component of the pro-active top-down selection mechanism (Desimone and Duncan, 1995; Nobre and Stokes, 2019). While WM plays an important role in the volitional control of attention, it can also influence attentional selection in an involuntary way. Specifically, a stimulus encoded and actively maintained in visual WM automatically attracts attention upon a subsequent presentation (review: Soto et al., 2008a). This effect has been revealed, first, by dot-probe experiments, demonstrating faster responses to probes presented at the location of the WM-maintained stimulus, than to probes following the unfamiliar stimulus (Downing, 2000). Second, by visual search experiments, showing that the search times are increased when a WM-maintained stimulus appears in the search array as a distractor (e.g., Olivers et al., 2006; Soto et al., 2005, Soto et al., 2007a, 2009). Third, by eye-tracking studies indicating that eye-movements are attracted by visual input matching the WM content (Hollingworth et al., 2013; Schneegans et al., 2014; Silvis and Van der Stigchel, 2014). Finally, by electrophysiological experiments, which revealed that the WM-maintained stimuli evoke the N2pc component, a classic index of covert attention shifts (Carlisle and Woodman, 2013; Kumar et al., 2009). Importantly, such a WM-based attention capture effect is not a form of priming, as it is observed only when a stimulus is actively maintained in WM, but not when it is merely seen; and is considered automatic and involuntary, as it occurs even when detrimental to the task performance (review: Olivers et al., 2011).

The automatic guidance of attention from WM has been so far demonstrated mainly with the use of simple stimuli, defined either by colour or orientation (review: Soto et al., 2008). Stimuli varying on a single dimension of one basic feature are generally most effective in guiding bottom-up attention, as they can be processed pre-attentively and result in a pop-out search (Wolfe & Horowitz, 2017). Therefore, a question arises whether also complex stimuli, whose identity is typically defined by multiple features and relations among them, can cause a similar WM-based attention guidance effect. The Downing’s (2000) dot-probe study revealed that images of faces, abstract geometric shapes, and drawings of real life objects captured attention when maintained in WM. However, subsequent experiments using visual search paradigms found no evidence of attention capture when drawings of real life objects (Houtkamp and Roelfsema, 2006) or complex artificial shapes (Downing and Dodds, 2004; Peters et al., 2009; Zhang et al., 2010) were maintained in WM.

In the light of such conflicting findings we investigated whether two types of complex, naturalistic stimuli - images of faces and houses - are prioritized by attention when maintained in visual WM. Further, using electrophysiological data we aimed to track the time-course of neuronal activity involved in matching the perceptual input and the WM representation. Faces and houses were chosen as both categories are defined by multiple features and thought to exhibit similar levels of complexity (Filliter et al., 2016). However, given the special status of faces in the human visual system, we hypothesized they might benefit from attentional and memory advantage over houses (Curby et al., 2009, Farah et al., 1998; Tsao and Livingstone, 2008).

The conducted experiment consisted of multiple blocks, in which a template stimulus (faces or houses) was either memorized for later recollection (WM condition) or merely seen without the need to memorize (exposure condition). Next, in each block participants performed a sequence of dot-probe trials, in which the template and a control stimulus were presented as task-irrelevant distractors (MacLeod et al., 1986). We hypothesized that in the WM condition the templates will involuntarily attract attention, as indicated by an RT effect and the N2 posterior contralateral (N2pc) ERP component, but we did not expect to observe an attention capture in the mere exposure condition.

## Methods

### Subjects

We analyzed data of 28 participants (18 females, mean age = 24.2, SD = 2.59 yrs., range: 19-28, 2 left-handed). They all declared normal or corrected-to-normal vision and no history of mental or neurological disorders. Data of 6 additional participants were collected, but they were excluded from the analysis: 4 participants due to the technical problems during an EEG recording procedure; 1 participant did not comply with the dot-probe task instruction; and 1 participant due to insufficient number of epochs remaining after EEG signal pre-processing (detailed criteria are described in the “EEG recording and analysis” section).

The study was conducted with the approval of the human ethics committee of the SWPS University of Social Sciences and Humanities (Warsaw, Poland). All subjects provided written informed consent and received monetary compensation for their time (100 PLN= c.a. 25 EUR).

### Stimuli

Two sets of stimuli were used. First, 60 pictures of faces with neutral expression (30 male, 30 female; all Caucasian) selected from the Karolinska Directed Emotional Faces stimulus set (KDEF; Lundqvist et al., 1998). From their original format, the face photographs were converted to grayscale using the Gnu Image Manipulation Program (GIMP; available at http://www.gimp.org/). Second, 60 pictures of houses from the DalHouses stimulus set (Filliter et al., 2018). All houses’ images were originally in grayscale, presented on a white background, thus no modifications were introduced. Identifiers of stimuli used in the present study can be found in the project description at OSF (https://osf.io/9rc4j/).

### Procedure

The experimental procedure was written in the Presentation software (Neurobehavioral Systems, Albany, CA, USA) and presented on a FlexScan EV-2450 (Hakusan, Ishikawa, Japan) screen through an Intel Core i3 computer. Participants were sat comfortably in a dimly lit room with a viewing distance of 57 cm, which was maintained by a chinrest.

The procedure started with a display providing subjects with the task instructions and information about the trial structure. The procedure consisted of two tasks: a working-memory (WM) task and a mere exposure task; and involved two stimuli types: faces and houses. Thus, there were four conditions: a face WM condition; a house WM condition; a face exposure condition; and a house exposure condition; which were presented to participants as separate blocks, the order of which was randomized. Each condition was further sub-divided into 32 memory or exposure blocks. For each memory/exposure block one template and one control stimulus were randomly chosen from the pool of all houses or faces. Additionally, in the face conditions the face stimuli were gender-matched, i.e. female and male template images were paired only with, respectively, female and male control images. In both, a face WM condition and a face exposure condition female faces were used in half of the blocks, and male faces in the other half. All stimuli were presented against a black background.

Each of those 32 blocks started with a central presentation of a template stimulus for 5000 ms (Fig. 1). The instruction - either “Memorize this picture“ (in the WM condition) or “Take a look at this picture” (in the exposure condition) - was displayed above the image. The face images subtended 7.4° x 10.0° of the visual angle, while house images varied in size and subtended from 6.1° to 9.3° × 7.8° of the visual angle. After the display of the template stimulus, a sequence of dot-probe trials was presented. Each dot-probe trial started with a fixation cross (subtending 0.9° x 0.9° of the visual angle) displayed in the centre of the screen. The fixation cross remained on-screen throughout the trial. After 1000 ms a pair of stimuli were presented bilaterally for 200 ms - the template stimulus on one side and control stimulus on the other. Face stimuli were presented with their inner edge 4.4° left and right from the fixation cross, while house stimuli with the inner edge from 3.8° to 5.3° left and right from the fixation cross. Next, a target asterisk subtending 0.7° x 0.7° of the visual angle was presented for 150 ms in the location of the centre of either the template stimulus (congruent trial) or the control one (incongruent trial). Participants were instructed to maintain their gaze on the centrally presented fixation cross, ignore the laterally appearing stimuli, and indicate the side of the target asterisk presentation (left or right) by pressing one of two buttons using index fingers of their left or right hand. Participants were asked to respond as quickly and accurately as possible. The response time to the target asterisk was limited to 3000 ms and the next trial started immediately after the manual response. Within each dot-probe sequence the template stimulus was presented on the left side in half of the dot-probe trials, and on the right side in the other half. Further, half of the dot-probe trials were congruent and half were incongruent. The order of trials within each sequence was randomized.

**Figure 1.**
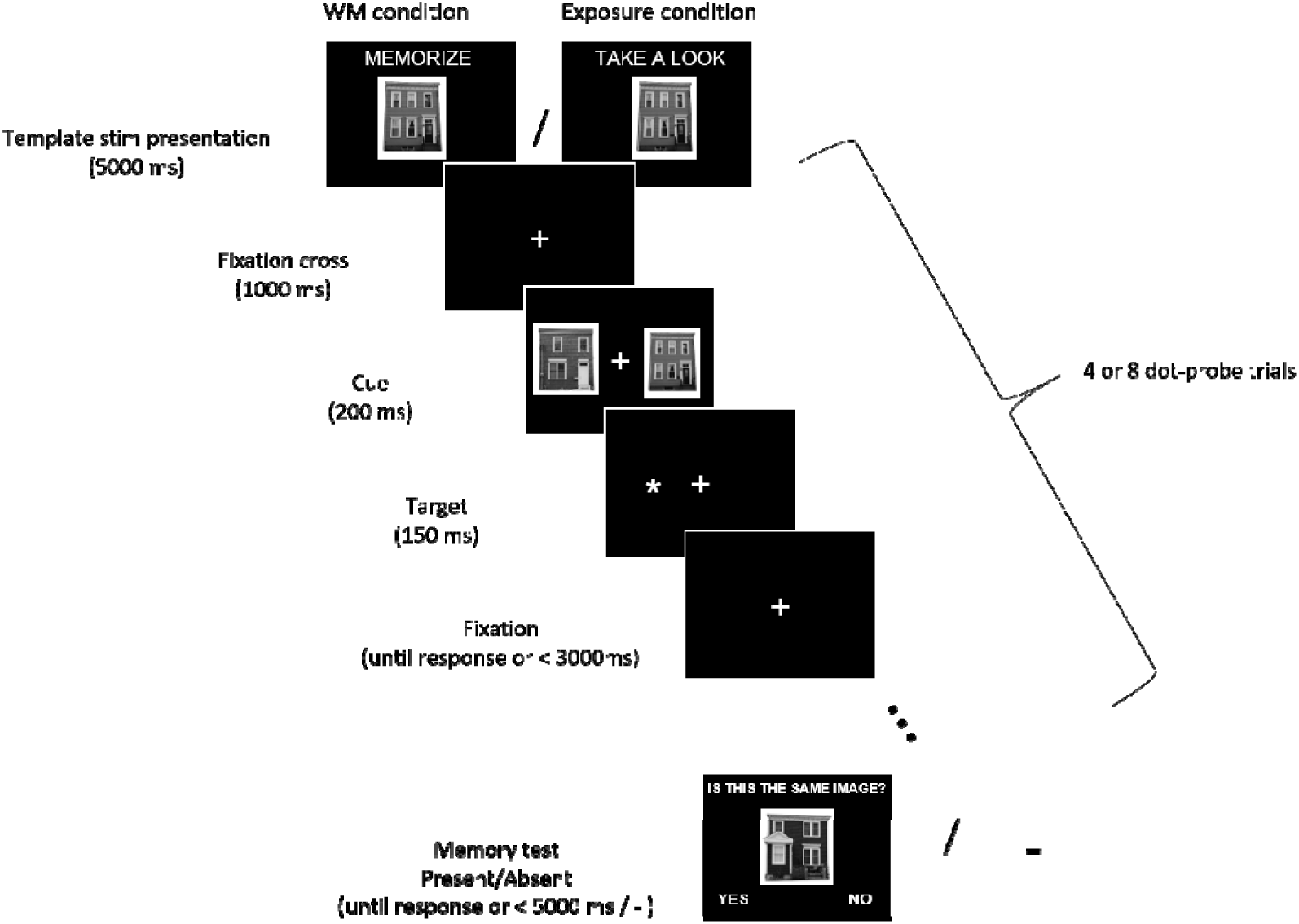
Experimental procedure. Participants were asked to memorize (WM condition) or just look (exposure condition) at the presented stimulus (either a face or a house). Then they performed 4 or 8 dot-probe trials in which their task was to indicate with a button press the location of a target asterisk (left or right). Asterisks were preceded by pairs of stimuli, which participants were supposed to ignore. A memorized/seen stimulus was presented on one side, and a control one on the other side. In the WM condition participants completed a memory test, in which they had to decide if the presented stimulus is the one they were instructed to memorize (in the exposure condition there was no memory test).

In the WM conditions, participants were given a memory test at the end of each memory block (i.e. after completing the dot-probe trials sequence). Either a template image or a different image (neither a template, nor a control stimulus) was presented centrally. Participants had to indicate whether the presented stimulus is the same or different from the one they were previously asked to memorize. Above the image a question “Is this the same image?“ was displayed. In half of the WM blocks the stimulus was the template image (correct answer “yes”), and in the other half it was a different image (correct answer “no”). The answer “yes” was displayed in the left corner of the screen and the answer “no” in the right corner. Participants responded by pressing one of two buttons (left for “yes”, right for “no”). The response time in the memory test was limited to 5000 ms and the next WM block started immediately after the manual response. In the mere exposure condition participants were not tested for stimulus recognition, but immediately after completion of the dot-probe sequence the next exposure block started.

In total 192 dot-probe trials were presented per condition. Within each condition half of the WM/exposure blocks comprised 4 dot-probe trials, and the other half comprised 8 dot-probe trials (the order of blocks was random). The number of trials varied in order to prevent participants from expecting the exact moment of a memory test, and thus encourage them to maintain the WM active throughout the block. Participants had a self-paced break five times per condition.

### Analysis of behavioral data

All analyses of behavioural data were conducted using custom-made Python scripts. Accuracy of responses to the presentation side of the target-dot was calculated as a percentage of correct responses. The obtained values are presented in the Results section, but due to ceiling level performance in the majority of participants this measure was not analysed statistically. Therefore, analysis of the dot-probe task data was focused on establishing whether reaction times (RT) of manual responses to the target asterisk differ between two types of trials: those in which the asterisk was presented on the same side as the potentially attention-grabbing stimulus (memorized/seen face or house; congruent trials) and those in which the asterisk was presented on the neutral stimulus side (control face or house; incongruent trials). Mean reaction times were calculated only for the correct responses. For the WM condition the accuracy of memorizing a template stimulus was calculated as a percentage of correct responses in the memory test.

### EEG recording and analysis

During the experiment, EEG signal was recorded with 64 Ag-AgCl electrically shielded electrodes mounted on an elastic cap (ActiCAP, Munich, Germany) and positioned according to the extended 10–20 system. Vertical (VEOG) and horizontal (HEOG) electrooculograms were recorded using bipolar electrodes placed at the supra- and sub-orbit of the right eye and at the external canthi. Electrode impedances were kept below 10 kΩ. The data were amplified using a 128-channel amplifier (QuickAmp, Brain Products, Enschede, Netherlands) and digitized with BrainVisionRecorder® software (Brain Products, Munich, Germany) at a 500 Hz sampling rate. The EEG signal was recorded against an average of all channels calculated by the amplifier hardware.

EEG and EOG data were analyzed using EEGlab 14 functions and Matlab 2016b. First, all signals were filtered using high-pass (0.5 Hz) and low-pass (45 Hz) Butterworth IIR filter (filter order = 2; Matlab functions: *butter* and *filtfilt*). Then data were re-referenced to the average of signals recorded from left and right earlobes, and down-sampled to 250 Hz. All data were divided into 768 dot-probe epochs (192 epochs per condition; [-200, 1200] ms with respect to the faces/houses images onset) and the epochs were baseline-corrected by subtracting the mean of the pre-stimulus period (i.e. [-200, 0 ms]). Further, epochs were rejected based on the following criteria (all values Mean±SEM): i) when there was no manual response to the target dots until 1.2 s after the onset (18.9±6.0; range [0, 120] epochs per subject); ii) when activity of the HEOG electrode in the time-window [-200, 500] ms exceeded −40 or 40 uV (105.5±19.1; range [11, 352] epochs per subject); iii) when activity of the P7 or P8 electrode in the time-window [-200, 600] ms exceeded −80 or 80 uV (none of the epochs rejected). Thus, after applying the described criteria the average number of analyzed epochs per subject was: 643.6±22.0; range [355, 752].

A subject was excluded if the number of epochs in any condition was < 60. This criterion resulted in excluding 1 subject out of 29 (but additional 5 subjects were excluded due to other criteria, as described in the “Subjects” section). The numbers of epochs provided above were calculated based on the final sample of 28 subjects.

Next, each EEG-EOG data-set was decomposed into 50 components using Independent Component Analysis as implemented in the EEGlab *pop_runica* function. To remove residual oculographic artefacts from the data the following procedure was used: time-course of each component was correlated with time-courses of HEOG and VEOG electrodes and in case the Spearman correlation coefficient exceeded −0.3 or 0.3 a component was subtracted from the data. Using this procedure 3.0±0.2 components (range [1, 6]) per subject were removed.

After applying the described preprocessing steps, data were divided with respect to the condition and presentation side of the template stimulus. The N2pc is calculated as the difference between the contralateral and ipsilateral waveforms. To create those waveforms we used the signal from P8 and P7 electrodes. Specifically, when template stimulus was presented on the left side, P8 was the contralateral electrode and P7 was the ipsilateral electrode. When template stimulus was presented on the right side, P7 was the contralateral electrode and P8 was the ipsilateral electrode. For each condition contralateral and ipsilateral signals were first concatenated and then averaged, resulting in the creation of contralateral and ipsilateral waveforms. The averaged waveforms within the 200-400 ms temporal window was used for the N2pc analysis, which is largely consistent with previous studies (e.g. Woodman et al., 2009; Reutter et al., 2017, Wójcik et al., 2019). Further, based on the visual inspection of the obtained ERP waveforms a 100-200 ms window was included in the factorial analysis.

### Statistical analysis

Statistical analyses were conducted in the JASP software and cross-checked with Statcheck (http://statcheck.io/index.php). The values are reported as Mean±SEM, unless stated otherwise.For all statistical tests probability values were reported (*p*) and the standard 0.05 alpha level was used as a threshold for refuting the null hypothesis.

To test for the presence of the behavioral (RT) and electrophysiological dot-probe task effects repeated - measures (rm) ANOVA models were used. Specific to the analysis of RT was the factor of *congruency*, defined by the asterisk presentation side with respect to the memorized/seen item (congruent vs. incongruent trials). Specific to the electrophysiological analysis was the factor of *side*, defined the side on which ERP activity was recorded, with respect to the memorized/seen item (ipsi- vs. contra-lateral). The *side* effect was analyzed separately for activity recorded in early (100-200 ms) and late (200-400 ms) time windows. The factors of a *stimulus* (faces vs. houses) and *task* (memory vs. mere exposure) were included in all models. The simple main effects analyses were also conducted. Results were reported as *F*(df) and partial eta-squared, the indicator of the effect size, was reported as *η*_*p*_^*2*^.

To compare the accuracy scores in the memory test between faces and houses conditions, the data distribution was first tested with the Shapiro-Wilk test and, as it deviated from normality, a nonparametric two-tailed Wilcoxon test was used. The statistic was reported as a sum of positive ranks (*W*), together with the matched rank biserial correlation (*r*_*rb*_) as a measure of the effect size.

## Data availability

The data used in the statistical analysis can be accessed from the OSF (https://osf.io/9rc4j/). Raw EEG data, and scripts used for analysis presentation of the procedure will be shared by authors per request.

## Results

### Memory accuracy

High working-memory (WM) accuracy scores were observed for both faces (97.2±0.7%) and houses (94.2±1.6%) indicate that participants were actively maintaining the template stimulus in working memory. Comparing the WM accuracy between face and houses we found better memory performance for face images (*W* = 172.500, *p* = 0.010, *r*_*rb*_ = −0.150), which is in line with previous studies (Curby et al., 2009). However, due to the ceiling level performance and low effect size any conclusions should be treated with caution.

### Dot-probe task - behavioral results

In the dot-probe task participants exhibited ceiling-level accuracy (i.e. in indicating the target-dot presentation side), with the percentage of correct responses being: 97±0.6% in the house WM condition; 97±0.6% in the face WM condition; 97±0.8% in the house exposure condition; 98±0.5% in the face exposure condition. Therefore, the reaction times (RT) of correct responses to the target asterisk were analyzed as an index of attention capture. Specifically, we investigated whether RT were shorter when the target followed a potentially attention-grabbing template stimulus (i.e. congruent trials), in comparison to trials when it followed a control stimulus (i.e. incongruent trials; Fig. 2). In a three-way rm-ANOVA analysis we found a significant main effect of *congruency* (*F*(1, 27) = 15.49, *p* < 0.001, *η*_*p*_^*2*^ = 0.365) and *task* (WM vs. exposure; *F*(1, 27) = 9.05, *p* = 0.006, *η*_*p*_^*2*^ = 0.251), and a significant interaction between those two factors (*F*(1, 27) = 14.24, *p* = 0.001, *η*_*p*_^*2*^ = 0.345). With regard to this interaction there was a significant simple main effect of *congruency* in the memory condition (congruent trials: 352.06 ±8.67 ms; incongruent 368.54±10.86 ms; *F*(1, 27) = 21.42, *p* < 0.001, *η*_*p*_^*2*^ = 0.442), but not in the exposure condition (congruent trials: 349.33±11.14 ms; incongruent 350.55 ±10.56 ms; *F*(1, 27) = 0.265, *p* = 0.611, *η*_*p*_^*2*^ = 0.010). The main effect of *stimulus* (*F*(1, 27) = 0.052, *p* = 0.822, *η*_*p*_^*2*^ = 0.002) and other interactions (*task* x *stimulus*: *F*(1, 27) = 0.830, *p* = 0.370, *η*_*p*_^*2*^ = 0.03, *stimulus* x *congruency: F*(1, 27) = 0.033, *p* = 0.858, *η*_*p*_^*2*^ = 0.001, *congruency* x *task* x *stimulus: F*(1, 27) = 0.12, *p* = 0.746, *η*_*p*_^*2*^ = 0.004) did not reach significance. Therefore, in line with our hypothesis, stimuli which were actively maintained in WM (but not those merely seen) induced the attentional bias, and this WM-based effect was observed irrespective of the stimulus type.

**Figure 2.**
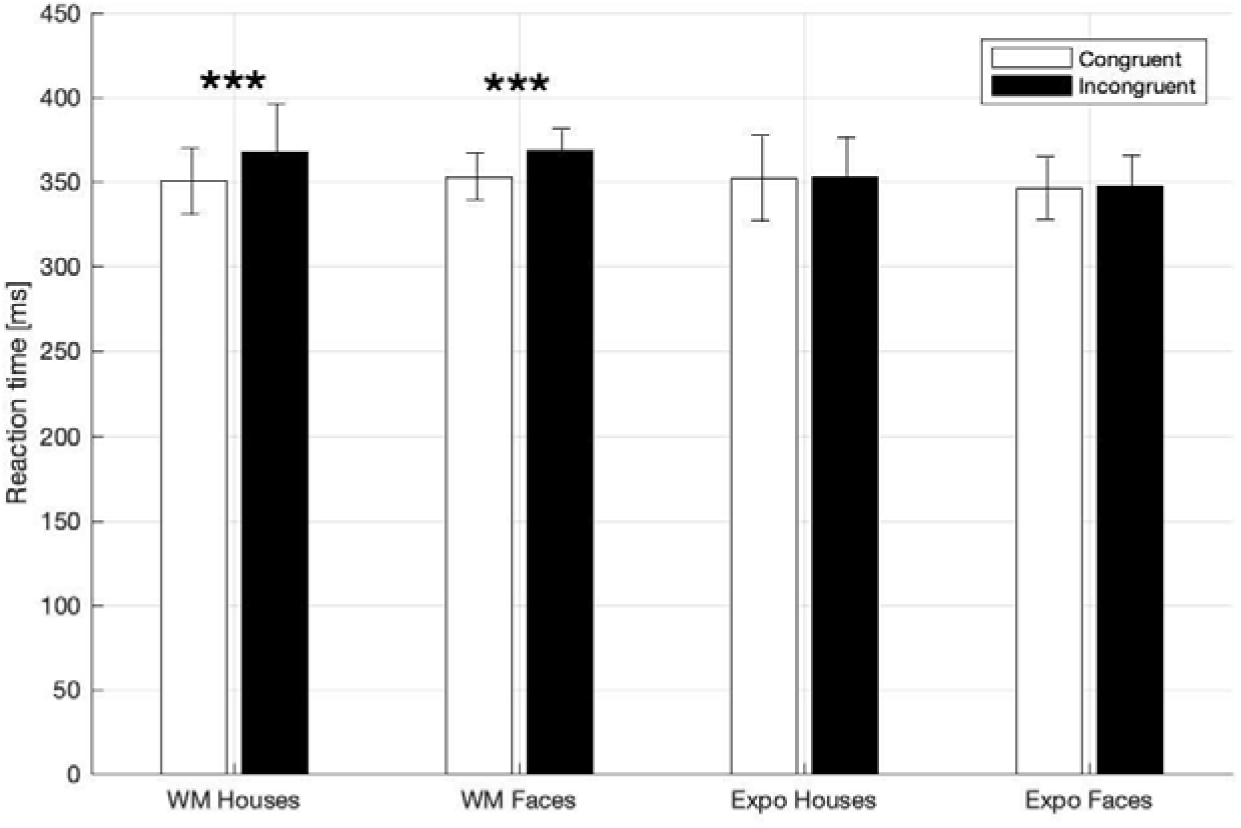
Mean RTs in dot-probe task for the WM and exposure conditions for both types of stimuli (House, Face). Asterisks indicate statistically significant difference between congruent and incongruent trials (*p* < 0.001). Error bars represent 2SEM.

### Dot-probe task - electrophysiological results

EEG data were collected to analyze the N2 posterior-contralateral (N2pc) ERP component, which is considered to be a robust index of covert attention shifts (Eimer, 1996; Kiss et al., 2008). N2pc is defined as a lower amplitude of an ERP recorded from the contra-lateral side with respect to the stimulus, in comparison to an ERP recorded ipsi-laterally. To calculate ipsi- and contralateral values we used signals from the P7/P8 electrodes averaged within the 200 - 400 ms time window (Fig. 3). The results of three-way rm-ANOVA (Table 1) indicate a significant *side* effect only (ipsilateral amplitude: 1.64±0.44 µV; contralateral: 1.49±0.44 µV). However given the lack of significant interactions between *side* and *task*, or *side* and *stimulus* we conclude that our manipulations did not affect the N2pc component. Thus, in contrast to our expectations, we did not observe larger N2pc in the WM task.

**Table 1.** Rm-ANOVA analysis of the electrophysiological effects. Analysis was conducted separately for 2 time windows (100-200 ms and 200-400 ms). Each model included the following 3 factors: *side* (recording from contralateral/ipsilateral electrodes), *task* (memory/exposure), *stimulus* (face/house)

| Factor | Time window:<br>100-200 ms |  |  | Time window:<br>200-400 ms |  |  |
| --- | --- | --- | --- | --- | --- | --- |
| | $F(1,27)$ | $p$ | $\eta_p^2$ | $F(1,27)$ | $p$ | $\eta_p^2$ |
| Side | 1.556 | 0.223 | 0.054 | 10.194 | 0.004 | 0.274 |
| Task | 0.661 | 0.423 | 0.024 | 0.032 | 0.860 | 0.001 |
| Stimulus | 31.910 | <.001 | 0.542 | 1.023 | 0.321 | 0.037 |
| Side x Task | 4.923 | 0.035 | 0.154 | 1.899 | 0.180 | 0.066 |
| Side x Stim | 0.066 | 0.321 | 0.036 | 0.069 | 0.795 | 0.003 |
| Task x Stim | 2.341 | 0.138 | 0.080 | 0.027 | 0.871 | 0.001 |
| Side x Task<br>x Stim | 4.923 | 0.035 | 0.154 | 2.032 | 0.165 | 0.070 |

**Figure 3.**
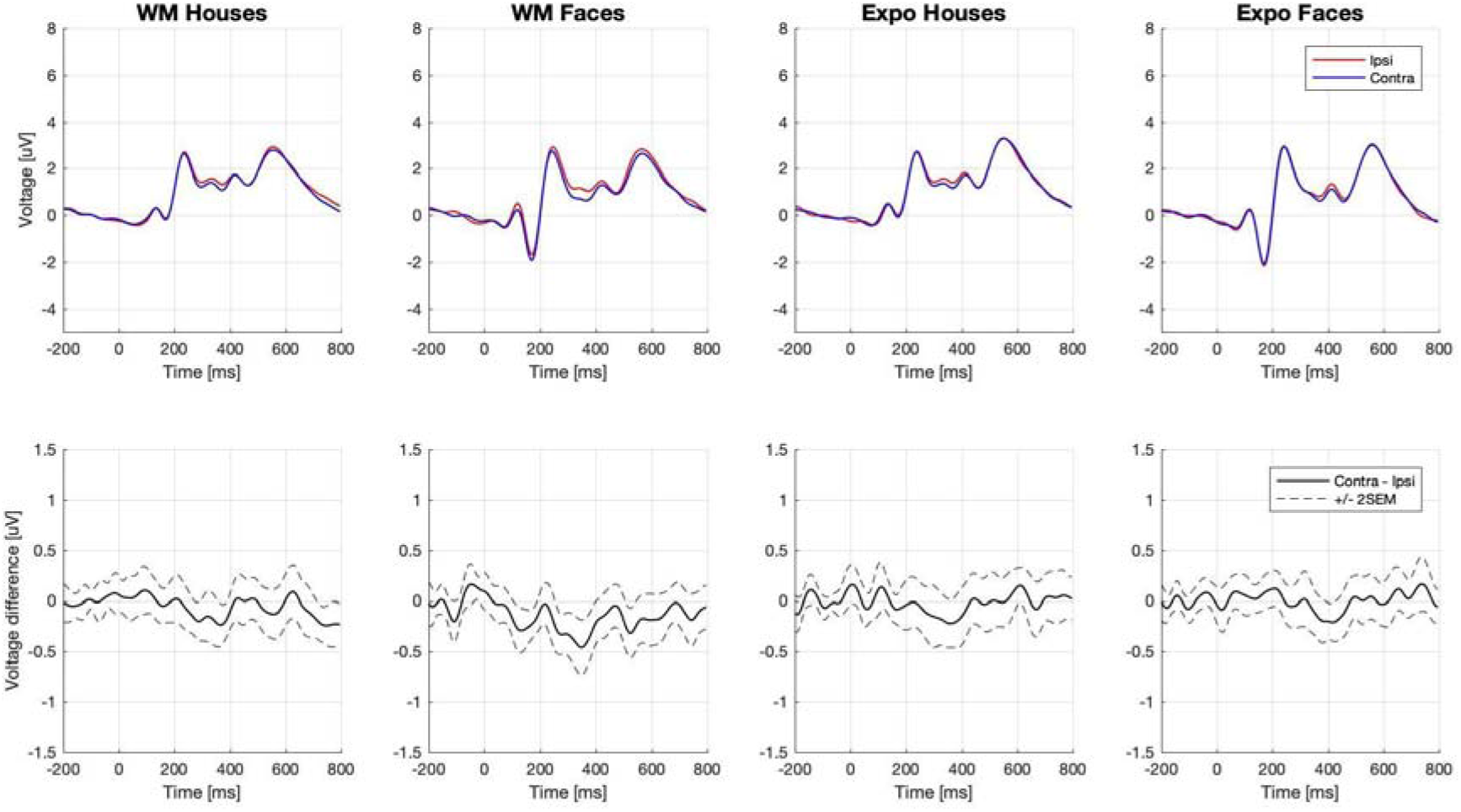
Event related potentials in the dot-probe task. Electrodes P7/P8 were chosen for the analysis. Waveforms recorded ipsi- and contra-laterally with respect to the seen or memorized stimulus are presented in the top row. Difference waveforms (i.e. contra - ipsi-lateral side) are presented in the bottom-row.

Inspection of the obtained ERP waveforms prompted us to conduct an exploratory analysis of lateralized activity in an earlier time-window (100-200 ms). This analysis yielded a significant interaction between *side*, *task*, and *stimulus* (Table 1). The simple main effects analysis showed that an early contralateral negativity can be observed in memory condition for faces (ipsilateral: - 0.51±0.39 µV; contralateral: −0.74±0.38 µV; *F*(1, 27) = 10.13, *p* = 0.004, *η*_*p*_^*2*^ = 0.273), but not for houses (ipsilateral: 0.15±0.43 µV; contralateral: 0.15±0.44 µV; *F*(1, 27) = 0.004, *p* = 0.951, *η*_*p*_^*2*^ < 0.001). In the mere exposure conditions the simple main effects were not significant neither for faces (ipsilateral: −0.85±0.42 µV; contralateral: −0.79±0.41µV; *F*(1, 27) = 1.02, *p* = 0.322, *η*_*p*_^*2*^ = 0.036) nor for houses (ipsilateral: 0.23± 0.42 µV; contralateral: 0.23±0.43µV; *F*(1, 27) = 0.01, *p* = 0.971, *η*_*p*_^*2*^ < 0.001). Therefore, as our analysis revealed, the very early lateralized activity was evoked only by the face images maintained in WM.

## Discussion

The present study examined whether complex naturalistic stimuli that are actively maintained in WM attract visual attention upon subsequent presentation. Such a WM-guided attentional selection has been consistently observed for simple stimuli (review: Soto et al., 2008a), but inconsistent findings were reported when more complex stimuli were used (Downing, 2000; Downing and Dodds, 2004; Houtkamp and Roelfsema, 2006; Peters et al., 2009; Zhang et al., 2010). Therefore, we conducted a study, in which images of faces and houses were either memorized or merely seen by participants, and subsequently presented in a dot-probe task. To test their capability to automatically attract attention we analyzed behavioral (RT) and electrophysiological (N2pc) indexes of attentional prioritization.

### Attentional prioritization of complex stimuli

Our main finding is that RTs were significantly shorter when the target asterisk followed a memorized face or house, in comparison to the situation when it followed a control stimulus, which indicates that the memorized stimuli were indeed prioritized by attention. Importantly, the RT effect was not observed, neither for faces nor for houses, when subjects merely saw the template images without an instruction to memorize them. Thus, first, our behavioural results replicate the findings of Downing (2000), who found a similar effect for complex stimuli in a dot-probe task. Second, they are in line with the wealth of visual search experiments showing increased RTs when a simple stimulus maintained in WM appeared in the search array as a distractor (Soto et al., 2005, 2007a, 2007b, 2008b, 2009; Olivers et al., 2006). Therefore, our study confirms and further extends the scope of the WM-based attention guidance effect and, in more general terms, provides evidence that complex naturalistic stimuli are able to guide attention in an automatic manner (see: Downing, 2000; Wolfe and Horowitz, 2017).

Importantly, in contrast to Downing’s (2000) and our results, several previous studies reported that complex but artificial shapes (Downing and Dodds, 2004; Peters et al., 2009; Zhang et al., 2010) or drawings of real-life objects (Houtkamp and Roelfsema, 2006) did not attract attention when maintained in WM. This led Zhang et al. (2010) to conclude that the attentional guidance critically depends on the stimulus features, with a stronger effect for simple than for complex stimuli. However, such conclusion disagrees with Downing’s (2000) and our studies, which show that even stimuli defined by multiple features and by relations among them can induce WM-based attentional biases. This indicates that the mere complexity of a stimulus is not likely to be a critical factor in the investigated phenomenon. We rather argue that our stimuli might have been processed holistically and perceived as more meaningful, in contrast to the artificial ones. This is in line with Xu (2017), arguing that visual WM is not typically used to encode features of a single dimension, but rather to store integrated representations of meaningful objects.

Two opposing accounts of the mechanism behind the WM-based guidance effect have been proposed. First one emphasizes the role of verbal (and perhaps semantic) representations in WM maintenance and subsequent directing of visual attention. It is based on studies showing that verbalization by itself can induce attention guidance (Soto and Humphreys, 2007a), and that even when visual stimuli are used as memory items the articular suppression task impairs the guidance effect (Soto and Humphreys, 2008b; Downing and Dodds, 2004; Woodman and Luck, 2007). However, the second view assumes that the guidance effect relies predominantly on visual representations. It is supported by experiments revealing the effect only when stimuli were defined by small and hard to verbalize differences in their attributes (e.g. hues of one color or slightly differing shapes) but not when easy to verbalize categorical differences were used (Olivers et al., 2006). Importantly, the stimuli used in our study were also difficult to verbalize and required maintaining a predominantly visual representation. Thus, our results provide further support for the second view.

Finally, while the WM-based attentional prioritization of faces observed in the Downing’s dot-probe study (2000) is replicated by our dot-probe experiment, his findings for drawings of real-life objects were not replicated subsequent visual search experiments (see: Houtkamp and Roelfsema, 2006). Therefore, the question arises whether dot-probe and visual search procedures differ in their sensitivity to detect WM-based attention bias. Given the scarcity of studies using complex stimuli, this question definitely needs further investigation. Nevertheless, our results obtained with the face and house images confirm that influences of WM on attention might play a role and be further investigated also in more ecological conditions.

### Attentional prioritization - capture or hold?

The majority of previous studies investigated the WM-guidance effect using behavioral methods, and thus relevant EEG or fMRI data are scarce. Thus, in the present study we collected electrophysiological data, with the main aim of evaluating the time-course of attentional prioritization. However, the N2pc ERP component - a classic index of the attentional capture (Eimer, 1996; Kiss et al., 2008) - was not affected by the memory manipulation, which is in disagreement with our hypothesis and with previous studies showing larger N2pc in similar WM tasks (however, using simple stimuli; Carlisle and Woodman, 2013; Kumar et al., 2009). Thus, in our study we found a robust behavioral (RT) effect of attentional engagement, but at the same time no electrophysiological effect (N2pc). The potential explanation of the dissociation between RT and N2pc is that the memorized stimuli did not automatically capture attention (thus no N2pc effect), but rather held and engaged attention for a longer time. What further supports this interpretation is that we observed an elongation of RT in the incongruent trials, rather than shortening of RT in the congruent ones - this is evident when the WM and exposure conditions are compared (Fig. 2).

Importantly, previous visual search studies including valid, neutral, and invalid conditions have provided conflicting results on the matter of capture versus hold. Some found longer RTs in invalid as relative to neutral trials, but no difference between valid and neutral condition (which would be indicative of an attention hold by memorized items; Soto et al., 2007b). However, others show both shorter RT in valid and longer RT in invalid trial, in comparison to the neutral ones (which would be indicative of both capture and hold; Soto et al., 2006). Thus, further studies are required to elucidate the precise mechanism of attentional prioritization of the WM-maintained items.

Importantly, attentional prioritization occurs when stimuli maintained in WM are subsequently presented as task-irrelevant, but it is even stronger when stimuli are task-relevant (Carlisle and Woodman, 2011a, 2011b, 2013). Task-relevance might be a factor defining the extent of capture and hold, and absence of the N2pc effect might stem from the fact that the memory items were task-irrelevant in our study. Thus, future studies will investigate the effect of naturalistic stimuli in situations when they are task-relevant, which would also more closely reflect daily life situations. Finally, even though the observed effect is interpreted as a hold rather than capture of attention, we argue that it is rather automatic and involuntary than strategic in its nature. This is suggested by brief presentation time of distractor stimuli (200ms), the minimal demands on WM for the memory test, and the fact that occurence of the observed WM-based attention capture effect was detrimental to the task performance (in line with Downing, 2000). Such interpretation is in agreement with previous studies using simple items as WM templates (Olivers et al., 2011).

### Early prioritization specific to faces

Even though we did not find the N2pc effect in the planned analysis, in an exploratory analysis we did find electrophysiological evidence suggesting a very early prioritization of the memorized faces. Specifically, we observed a contralateral negativity in response to the memorized face already between 100 and 200 ms after the stimulus onset (i.e. co-occurring with the P100 component). Thus, it is not clear whether contralateral negativity occurring so early can be termed N2pc, as N2pc is considered to occur around 175-200 ms after the stimulus onset (i.e. co-occurring with N2; e.g. Woodman et al., 2009; Reutter et al., 2017, Wójcik et al., 2019).

Observing contralateral negativity already around 100 ms after the stimulus suggests it represents the early and perceptual stages of processing. Enhanced activity of the occipital area in response to a stimulus held in WM has been already reported (Tan et al., 2014, 2015). The difference is that Tan and colleagues analyzed P100 amplitude, and here we analyzed contralateral negativity (i.e. in our study the WM-maintained stimulus was presented always in pair with control stimulus, thus analysis of P100 is not possible). However, others did not observe any evidence for such an early WM-associated activity (e.g. Kumar et al., 2009; Telling et al., 2010). It is thus important to emphasize that in our study the early effect was present for faces, but not for houses. While the mechanisms of such early electrophysiological effect remain to be investigated, the fact that in our study it was observed for faces is in line with several lines of evidence. First, due to their evolutionary and social importance, faces are processed in a largely automatic and holistic manner (Farah et al., 1998; Tsao and Livingstone, 2008). Second, due to holistic encoding strategies, faces benefit from a WM advantage (which is observed also in our data; Curby et al., 2009). Third, continuous flash suppression (CSF) studies show that faces actively maintained in WM break the CFS faster than faces that were merely seen (Gayet et al., 2013; Pan et al., 2014). The fact that in the CFS paradigm WM can bias face perception outside of awareness is in line with the automatic and involuntary (possibly pre-attentive) nature of the effect found here. Thus, the attentional prioritization revealed in the present study is a plausible mechanism accounting for the CFS effects. Finally, face recognition is performed by a specialized set of brain regions (Haxby et al., 2000; Kanwisher and Yovel, 2006) with the initial stages of face categorization occurring as early as 80-150 ms post-stimulus (Herrmann et al., 2005), which is in line with the observed early effect.

The presence (or absence) of early occipital cortex activity in response to the WM-maintained stimuli is relevant to the ongoing debate on the neuronal mechanisms of visual working memory. Here, two opposing theories have been proposed: first, the top-down amplification hypothesis, which assumes that visual WM items are maintained by fronto-parietal interactions (Bettencourt and Xu, 2016; Christophel et al., 2018, Riley and Constantinidis, 2016, Thigpen et al., 2019; review: Xu et al., 2017); second, the sensory-recruitment hypothesis, assuming that visual WM items are stored and maintained in the visual cortex (i.e. that perception and visual WM share the same neural substrate; Postle, 2006). The latter view might particularly effectively account for the automatic interactions between perception and the WM contents, which were observed in our and other studies (e.g., Albers et al., 2013, Gayet et al., 2017, Silvanto and Cattaneo, 2010, Teng and Kravitz, 2019). Importantly, given that ERP components observed in the 100-200 ms time-range are generated by sensory brain regions and reflect perceptual processing (Nusslock, 2016), such an early prioritization of the memorized faces provides support for the sensory recruitment theory. Further, such an early effect was not found in previous dot-probe studies using very salient and relevant emotional faces (Holmes et al., 2009) or self-faces of participants (Wójcik et al., 2019; Bola et al., 2020), which further indicates it might specifically reflect a match between the WM-maintained representation and an incoming stimulus. However, because this analysis was exploratory, the conclusion should be treated with caution.

In conclusion, by providing evidence that complex naturalistic stimuli maintained in WM are prioritized by attention our study replicates and extends the investigated effect. Our results are thus consonant with emerging evidence for the prospective and dynamic role of memory in guiding perception and behaviour. Further, we provide evidence for a very early prioritization of the memorized face images, which resonates well with previous studies indicating preferential processing of faces. Finally, the present study encourages further investigation of the WM-based influences on attention in more ecological settings.

## Notes

**Conflict of interest:** The authors declare no competing interests.

**Funding:** This study was supported by a National Science Center Poland (grant number 2018/29/B/HS6/02152).

https://osf.io/9rc4j/

### Competing Interest Statement

The authors have declared no competing interest.

